# A framework for offline evaluation and optimization of real-time algorithms for use in neurofeedback, demonstrated on an instantaneous proxy for correlations

**DOI:** 10.1101/351072

**Authors:** Michal Ramot, Javier Gonzalez-Castillo

## Abstract

Interest in real-time fMRI neurofeedback has grown exponentially over the past few years, both for use as a basic science research tool, and as part of the search for novel clinical interventions for neurological and psychiatric illnesses. In order to expand the range of questions which can be addressed with this tool however, new neurofeedback methods must be developed, going beyond feedback of activations in a single region. These new methods, several of which have already been proposed, are by their nature complex, involving many possible parameters. Here we suggest a framework for evaluating and optimizing algorithms for use in a real-time setting, before beginning the neurofeedback experiment, by offline simulations of algorithm output using a previously collected dataset. We demonstrate the application of this framework on the instantaneous proxy for correlations which we developed for training connectivity between different network nodes, identify the optimal parameters for use with this algorithm, and compare it to more traditional correlation methods. We also examine the effects of advanced imaging techniques, such as multi-echo acquisition, and the integration of these into the realtime processing stream.

## Introduction

The field of real-time fMRI based neurofeedback is growing rapidly, prompting the equally rapid development of new methods and algorithms designed to make this tool ever more flexible (1–6). A particularly interesting new method is connectivity-based neurofeedback, in which the goal is to feedback correlations between different brain regions, rather than the activity levels in a single region. From a basic science perspective, this approach is particularly appealing because it opens up an entirely new space of potential targets for neurofeedback, and moves away from the obvious over-simplification of a single brain region working in isolation. It allows us to investigate networks, and how interactions within and between networks affect behavior (7).

This approach to neurofeedback also holds great promise from a clinical perspective, as aberrant connectivity has been demonstrated in many neuropathologies, and is often the network characteristic most well correlated with symptom severity. In depression, studies have shown altered thalamo-cortical correlations (8), decreased functional connectivity within the reward network (9), and decreased cortico-cortico correlations (10). Aberrant connectivity has also been implicated in schizophrenia (11–14), ADHD (15, 16) and Autism Spectrum Disorders (17–20), among others.

However, there are several significant hurdles for carrying out real-time fMRI connectivity-based (or correlation-based) neurofeedback. The first stems from the inherently slow nature of the BOLD signal. This is a limitation above and beyond the slow acquisition rate, and is far more fundamental in that while the slow acquisition rate can be improved upon, the Hemodynamic Response Function (HRF) cannot be bypassed. The HRF not only imposes a delay between the neural event and the feedback, but it also limits the effective acquisition rate, as any data acquired with a TR of <1s will have substantial autocorrelations between neighboring time points (21). Secondly, the real-time fMRI signal is noisy, and only a limited set of preprocessing steps can be carried out in real-time. Thirdly, global signal fluctuations, comprised both of global artifacts such as motion and physiological noise, as well as real global signal, are prominent in fMRI data (22–25). These global fluctuations must be accounted for, to ensure that the correlation being reinforced is specific to the selected targets, rather than a global event.

In order to overcome these difficulties, we developed an instantaneous proxy for correlations, described in (26), to be used for correlation-based training. Unlike the block design or sliding window based correlational methods that have mostly been used previously for providing correlation-based feedback (27–29), this method allows for far more feedback events (relative to block design), and for much faster, less history-dependent feedback (relative to sliding windows). However, the development of this method raised many additional questions, most importantly relating to the validation and optimization of the chosen algorithm. Neurofeedback experiments are often lengthy and expensive, making it impractical to run several pilot studies to evaluate different potential algorithms. It is therefore important to assess the quality of correlational based methods / proxies, or indeed any other proposed algorithm, by offline simulations, prior to beginning any new neurofeedback experiment. Offline simulations (run on previously collected data), cannot of course be used to determine which algorithm or method leads to better or more robust learning. However, they can be used to compare the algorithm output to the “gold standard” calculations obtained by means of offline complex preprocessing of full series data (referred to in the rest of this manuscript as “fully processed“). This allows estimation of the effects of noise and timing present in the real-time signal. In this manner, previously acquired resting data can be used to optimize neurofeedback parameters such as target size, number of target regions, and their location, among other things.

We propose here a general framework for offline evaluation and optimization of candidate algorithms which can be used for any neurofeedback implementation, by using previously collected resting state data. In this framework, both the fully preprocessed data (i.e. the gold standard of offline processing, using the full dataset), and minimally preprocessed data, meaning data which is preprocessed only to the level that would be available to the real-time algorithm (assuming that the full time series is not available in a real-time environment) are run through the real-time algorithm, simulating the “real-time” algorithm output. During the evaluation phase, the algorithm output is compared to the relevant gold standard calculation carried out on the offline, full-series, fully processed data (Figure 1). This is done for both the fully processed data, to assess how the algorithm compares with full-series data, and with the minimally processed data, to assess how the algorithm fares given the noise levels in the real-time data.

**Figure 1.**
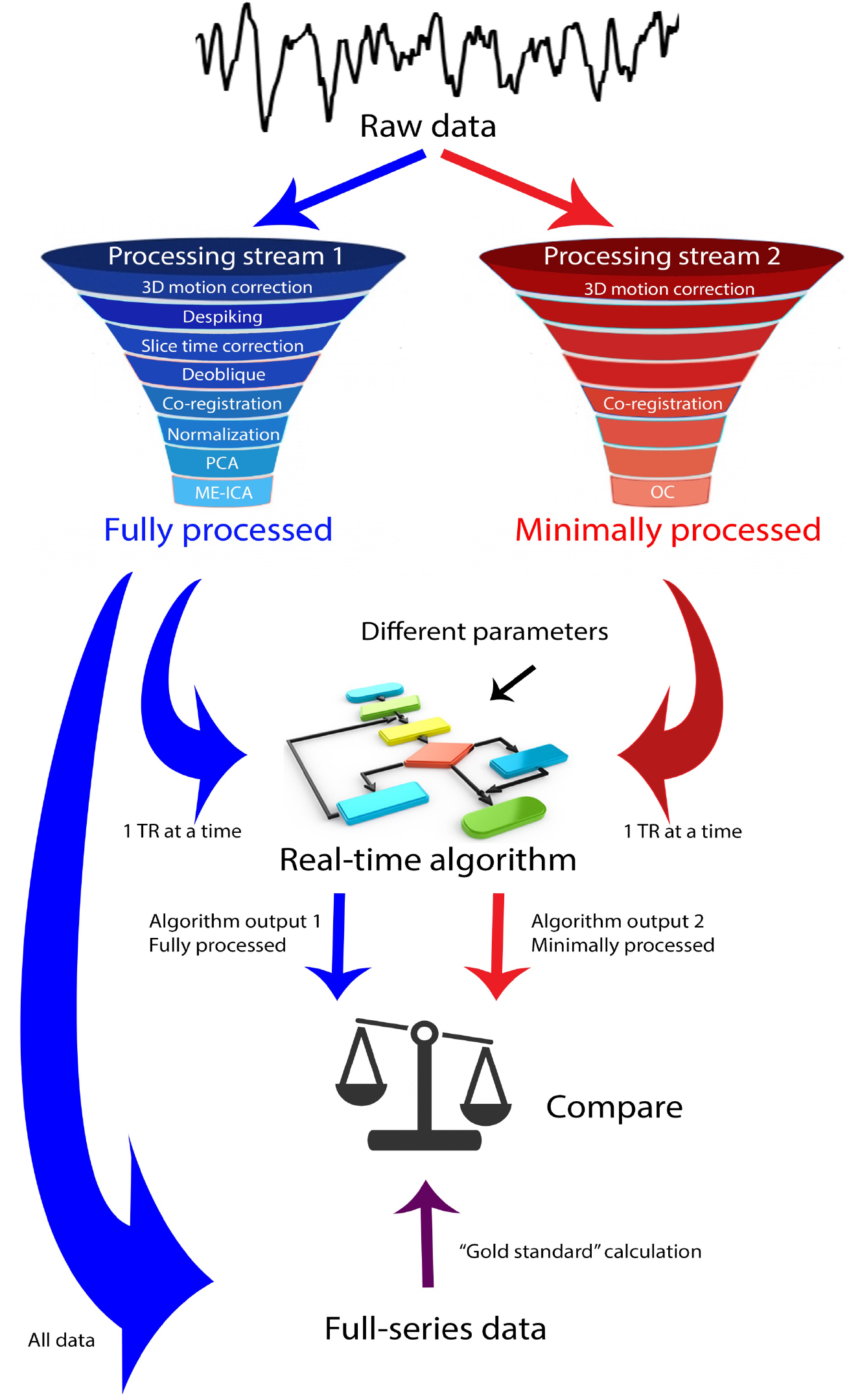
Framework for evaluation of candidate algorithms. Schema for evaluating potential real-time algorithms and optimizing their parameters using previously collected data. Raw data are processed in two processing streams, to output both fully processed data, which are the cleanest signals we can achieve, and minimally processed data, which represent the data processing steps which can be done online in realtime, and which the algorithm will have access to in a true real-time environment. Data from both processing streams are run through the algorithm, which goes through the same calculations it would perform in a real-time experiment. This results in two outputs, for each of the processing streams. The full series, fully processed data is also run through the “gold standard” calculation which would have been carried out offline, post-hoc. The output of this “gold standard” calculation and the algorithm outputs are then compared. Different algorithm parameters can then be evaluated, to uncover the optimal parameters in terms of correspondence to the gold standard calculation.

The gold standard calculation which the algorithm output is compared to can vary according to the goal of the training. We demonstrate the application of this framework on the two-point algorithm described in (26) and adapted for Figure 2, and explain how the many possible parameters are optimized. We also use this framework to compare the two-point method with more traditional correlation methods. Finally, we draw some general conclusions about real-time fMRI data, and optimization of this signal, for instance using multi-echo acquisition.

**Figure 2.**
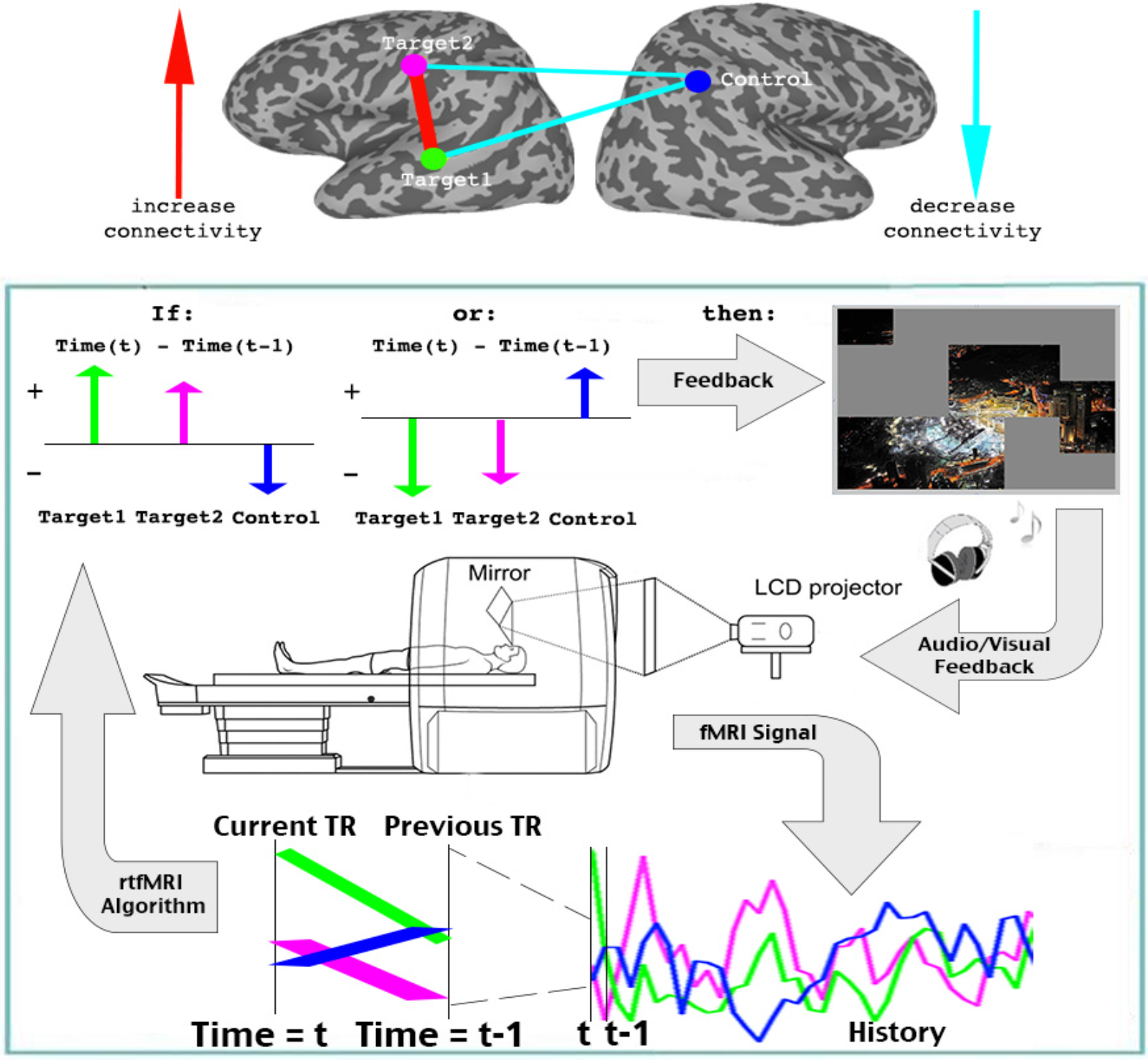
Two point algorithm. Each TR, the decision whether a feedback event occurs is made based on the change in signal from the previous time point (t-1) to the current time point (t) in the three ROIs (two targets and one control). Target locations are illustrations, not the targets used in the analysis here. A feedback event occurs if the direction of change in the two targets is the same, and opposite from the direction of change in the control ROI. Figure reproduced with minor changes from (26).

## Materials and Methods

### Participants

For the dataset used in these analyses, 34 typically developing young adults (20 female 14 male, mean age 23.4), with no history of psychiatric, neurological illness or other medical condition which could compromise cognitive function, were recruited. Participants were all right-handed, and all had normal or corrected to normal vision. Participants were all compensated for their participation, and gave written informed consent. The experiment was approved by the NIMH Institutional Review Board (protocol 10-M-0027, clinical trial number NCT01031407).

### Functional scans

All participants completed two 9-minute rest scans, for which they were instructed to lay still, not fall asleep, and fixate on the black fixation cross on a blank grey screen. Following the rest scans, participants completed two face/scene localizer scans, each 8 minutes twenty seconds long. The localizer scans were comprised of an initial 20 second blank grey screen with a fixation cross, and then sixteen 20-second long presentation blocks, each followed by 10 seconds of a blank grey screen with a fixation cross. During the presentation blocks, 20 pictures of either faces (face blocks) or scenes (scene blocks) were presented (stimulus duration = 200ms, inter-stimulus interval=700ms), with one or two of the images repeating in immediate succession in each block. To ensure attention to the images, subjects were given a repetition detection (1-back) task, and were instructed to search for repetitions, and to press a button on the response box whenever one occurred. There were 8 face blocks and 8 scene blocks in each of the localizer runs, and participants viewed 320 exemplars from each category, with each exemplar repeating no more than twice in each run.

### Imaging data collection and MRI parameters

All scans were collected at the Functional Magnetic Resonance Imaging Core Facility on a 32 channel coil GE 3T (GE MR-750 3.0T) magnet and receive-only head coil, with online slice time correction and motion correction. The scans included a 5 minute structural scan (MPRAGE) for anatomical co-registration, which had the following parameters: TE = 2.7, Flip Angle = 12, Bandwidth = 244.141, FOV = 30 (256 × 256), Slice Thickness = 1.2, axial slices. EPI was conducted with the following parameters: TR = 2s, Voxel size 3*3*3, Flip Angle: 60, Multi-echo slice acquisition with three echoes, TE_1_ = 17.5ms, TE_2_ = 35.3ms, TE_3_ =53.1ms, Matrix = 72×72, Slices: 28. 270 TRs were collected for the rest scans, and 250 TRs for the localizer scans. All scans used an accelerated acquisition (GE’s ASSET) with a factor of 2 in order to prevent gradient overheating.

### Definition of Regions of Interest (ROIs)

The localizer data was used to define individual face and scene ROIs for each participant. A standard General Linear Model was used with a 20 second long boxcar function, coinciding with the presentation blocks. This was convolved with a canonical hemodynamic response function, and deconvolved using the AFNI (Analysis of Functional Neuro-Images, (30)) function 3dDeconvolve. Face and scene selective ROIs were found using the face>scene contrast, or the scene>faces contrasts respectively. The functional and anatomical datasets were co-registered using AFNI, and then transformed to Talairach space. All ROIs, for each individual participant, were defined in Talairach space. In the faces>scenes contrast, we identified the center of mass for bilateral fusiform face area (FFA), occipital face area (OFA) and amygdala, as well as the right Anterior Temporal Lobe (ATL) face patch. In the scenes>faces contrast, we identified the center of mass of the bilateral Parahippocampal place area (PPA), and the bilateral dorsal scene patches. We then defined a spherical ROI of 4mm radius around each of these 11 centers of mass, to obtain 11 individually localized visual ROIs. These visual ROIs were used as the targets in all subsequent analyses, with all possible combinations of ROI pairs producing 55 unique target pairs ((11*11−11)/2 = 55). To define the control ROI, which was selected for being uncorrelated to the visual ROIs, we found the voxels least correlated to the visual ROIs across the entire group (N=34), and then defined a 4mm radius spherical ROI as the control. The control ROI was the same for all participants, for all target pairs.

### Real-time fMRI algorithm

The basic real-time algorithm used in these simulations is the same as that described in (26). Unless explicitly mentioned in the text, the algorithm gets input of the mean signal from two target ROIs and one control ROI. The objective is to selectively increase connectivity between the target ROIs, while simultaneously decoupling the targets from the control ROI, by providing feedback whenever this desired brain state is achieved. Each TR, the algorithm makes a decision whether feedback should be given (constituting a feedback event) based on whether or not the trend of the ROI mean signal has the same sign (increasing /decreasing) for the target ROIs, and the opposite sign for the control, between two time points, the current TR and the previous one. Mathematically, a positive feedback event for TR=t will happen if and only if:

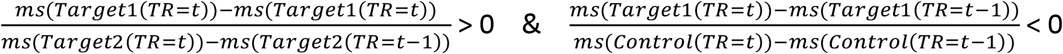

(ms = mean signal across all voxels in a given ROI)

If these two conditions are not met, then no feedback event occurs (see also Figure 2). This decoupling of the targets from the control ROI serves a dual purpose - first, it allows for the more selective reinforcement of target-target correlations, while avoiding feedback for global changes in activation. Such global fluctuations would be observed as changes in the momentary correlation measured by the algorithm between the two targets, but would also be seen as increased momentary correlation of the targets with control, i.e. the trend in all three ROIs would be in the same direction, and therefore no feedback would be given. It can also be useful to train more complex networks, where there is under-connectivity between some nodes and overconnectivity with others, as was the case in the original publication.

Our algorithm can also easily be extended to consider more than two adjacent time points. For example, the manipulation should the algorithm be using three time points for the decision, would look like this:

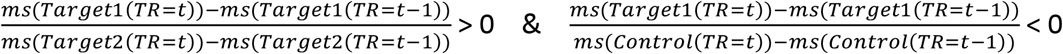

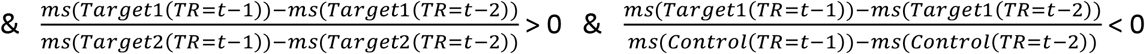

This means that the trend is required to be the same for the targets and opposite for the control both between the current TR and the previous one, and between the previous TR and the one before, for a positive feedback event to occur.

Similarly, the algorithm can be modified to accommodate the modulation of connectivity between more than two ROIs (or networks). For example, if interested in modulating the connectivity between three target ROIs, then a positive feedback event should only occur if the trend in all three targets has the same sign, and opposite from control; as in:

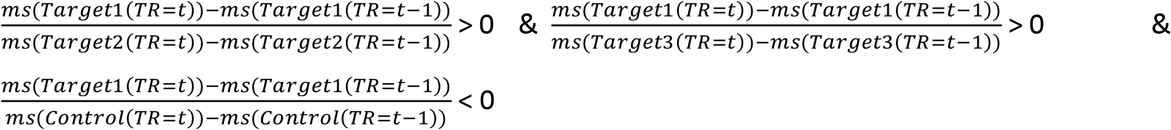

For four targets, the algorithm was more lenient, and required that only three of the four targets move in the same direction, for any given TR. This leniency was added because using the stricter requirement for all targets having the same trend resulted in very few feedback events. This configuration created 4 possible conditions for feedback, with three conditions in each one:

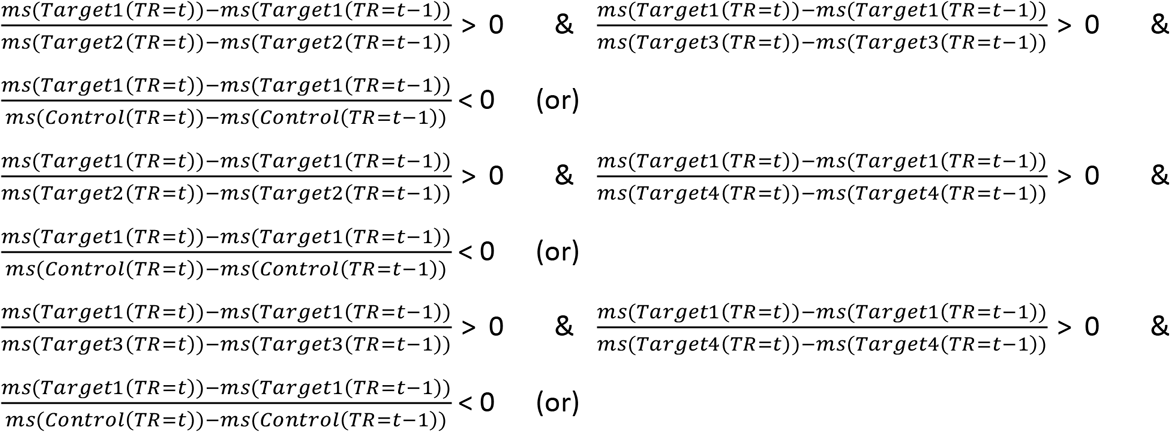

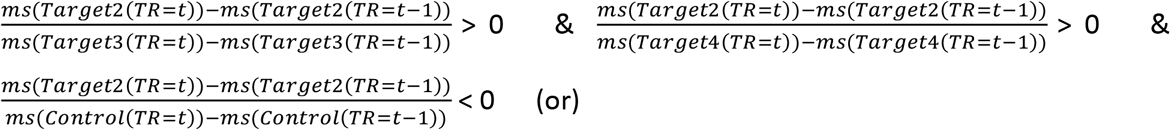

(ms = mean signal)

Another parameter to be manipulated and evaluated here, is ROI size. For our evaluation of the effect of ROI size, the algorithm remained the same as the basic algorithm, but the input changed so that it came from larger ROIs (defined using 6mm / 8mm / 10mm radius spheres centered around the same voxels identified as the center of mass for each region). The same is true for the evaluation of linear weighted combination of multi-echo data. In this case, instead of simply using as input to the neurofeedback algorithm the time series from the second echo (TE=35.3ms), we used as input the linear weighted combination of all echoes according to (31, 32). Such linear combination scheme, referred to as “optimally combined” (OC) in the rest of the manuscript, creates a new time series optimized for functional contrast (31, 32).

### fMRI offline data preprocessing

Post-hoc signal preprocessing was conducted in AFNI. For the fully processed data, which was processed using the gold standard for offline preprocessing, the first four EPI volumes from each run were removed to ensure remaining volumes were at magnetization steady state, and remaining large transients were removed through a squashing function (AFNI’s *3dDespike*). Volumes were slice-time corrected and motion parameters were estimated with rigid body transformations (through AFNI’s *3dVolreg* function). Volumes were co-registered to the anatomical scan. The data were then entered to a Multi-Echo ICA analysis (ME-ICA), as described in (33), to further remove nuisance signals (e.g., hardware-induced artifacts, residual head motion). Briefly, this procedure removes non-BOLD fluctuations (noise) present in the data based on the fact that BOLD and non-BOLD fluctuations differ in their properties across echo times, with signal from BOLD sources increasing linearly across echo times, and signal from non-BOLD sources remaining unchanged across echoes.

For the minimally processed data, the first four EPI volumes were also discarded, and then the volumes from all three echoes were corrected for motion using AFNI’s *3dVolreg* function. Volumes were co-registered to the anatomical scan. Unless otherwise stated, the minimally processed data used for the simulations refers to the second echo, which is similar in terms of echo time to a standard single echo acquisition (TE=35.3ms). An additional optional step, was to linearly combine the multi-echo data to optimize for functional contrast. The weights used for this purpose are given by

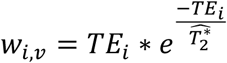

where *i = 1․.3* refers to echo, *v* refers to voxel, and 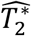 corresponds to voxel-wise estimates of 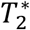 obtained via a log-linear firs to the multi-echo dataset. This data, comprised of the weighted average of the three echoes, was then used for all the simulations with optimally combined (OC) minimally processed data.

### Data analysis

All data were analyzed with in-house software written in Matlab, as well as the AFNI software package (30). Our “gold standard” calculation was designed to capture the same network being trained by the algorithm, so is defined as the correlation between the two targets, minus the average target-control correlation. It was computed by subtracting the average correlation of the two target/control pairs, from the target/target correlation:

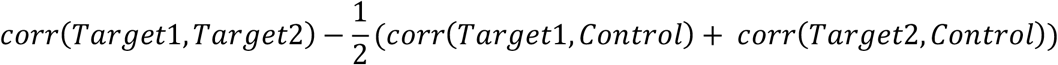

For the three-target and four-targets implementation of the algorithm, the composite measure was updated to reflect the greater number of targets. For three targets, it was defined thus:

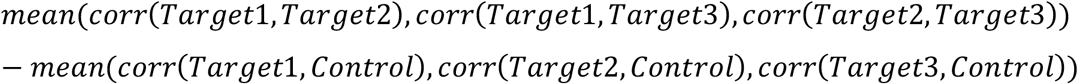

For four targets, it was the same, the average of all the pairs within the four targets, minus the average of the target-control pairs.

### Temporal Signal-to-Noise Ratio (tSNR)

The quality of each individual dataset was characterized in terms of its temporal Signal-to-Noise Ratio (tSNR). This metric is defined and computed on a voxel-wise basis as the ratio of the mean steady-state signal of the fMRI time-series to the voxel temporal standard deviation (34). We then averaged the tSNR across all voxels belonging to the two targets to get per-ROI tSNR values.

### Permutation Tests for Statistical Thresholding

All p-values were computed through permutation tests, randomly permuting the appropriate variables for 10,000 iterations. For instance, to calculate the p-value of the difference between two time points and three time points across the 55 target pairs, the real difference of the average for all 55 target pairs between the two-point and the three-point methods was compared to the distribution of a permutation test in which the two-point and three-point labels were permuted for each pair before averaging, and then the difference of the average of the permuted groups was calculated. This was repeated for 10,000 iterations.

## Results

First, we used the proposed evaluation framework to further validate the two-point method presented in (26), Figure 2 figure supplement 2, and test its generalizability. To that end, we tested the algorithm under different parameter configurations - number of time-points, number of ROIs, ROI size, using online optimal combination of echoes (OC) - on a different dataset than that used in the original publication. Apart from allowing us to validate the findings from (26) on an additional dataset, this dataset also had the advantage of having more participants (N=34); multiple individual functional Regions of Interest (ROIs) defined through a separate visual face/scene localizer; and being a multi-echo dataset it gave us the opportunity to test the effects of using the optimal combination of the echoes as explained in (33) as an additional preprocessing step in real-time.

Briefly, the algorithm has two possible outputs, reward / no reward, and acts as an instantaneous proxy for correlations between the defined target ROIs, and control ROI, using only two time points. Feedback events occur when the instantaneous change in the time-series (over the previous TR) is in the same direction for the target ROIs, and opposite in the control ROI ( see Materials and Methods, Figure 2).

The goal of this proxy is to train relative correlations between different brain regions, strengthening the correlations between the targets, while simultaneously decoupling them from the control region. Our “gold standard” calculation should capture the same network being trained, so is defined as the correlation between the two targets, minus the average target control correlation. This is calculated over the full time-series, using the fully processed data. We named this measure the composite difference correlation measure (hence referred to as the composite measure) (see Materials and Methods). The evaluation of the real-time algorithm is through its correspondence to this composite measure (Figure 1). To separate the effects of the online optimal combination of echoes, we first fed the algorithm data from the second echo only, which is comparable to single-echo data given its proximity to the average T2* of grey matter at 3T. Unless otherwise stated, results are based on this second echo.

### Number of time points

When designing an instantaneous proxy for correlation-based training, there are many factors to consider. While eventually we would want to test the performance of the algorithm on the minimally processed data that is actually available in the real-time setting, we must first begin by assessing how similar this instantaneous proxy is to the traditional, full-series Pearson’s correlations, and how that correspondence can be optimized. We began therefore by running the fully processed data through various implementations of the algorithm.

We first tested whether using more than two consecutive time points for the feedback would increase reliability in terms of correlation to the full-series data. To this end, we compared the algorithm output using two of our visual face targets (right fusiform face area, FFA and right occipital face area, OFA) and a control region identified as being uncorrelated to the targets (left inferior parietal lobule, IPL), for different configurations in terms of the number of time points the algorithm takes into consideration.

We first required the trend in the targets between two time points to be in the same direction and opposite from control, as in the original publication (Materials and Methods). This was calculated for each subject, for each of the two rest runs, and the total number of simulated feedback events was counted for each run. This was then compared with the composite measure described above. These data are shown in Figure 3a.

We then ran another set of simulations, in which the algorithm required the trend in the targets to be the same over three time points (see Materials and Methods). Once again, the number of simulated feedback events was recorded for each run, and compared with the composite measure (see Figure 3b).

As can be seen, the correlation of the two-point measure to the full-series composite measure is higher than that of the three-point method (r=0.65 vs. r=0.59). Using four time points lowered the correlations even further (r=0.34).

We next carried out the same analysis, using other target pairs. There were 11 visual targets, comprised of seven face regions and four scene related regions (see Materials and Methods). All regions were individually localized. While the control region was kept constant, as it was uncorrelated to all these visual areas, all 55 possible target pair combinations were examined. For each pair of targets, we calculated simulated feedback events using both the two-point measure and the three-point measure across all 34 subjects and both rest scans, and then calculated the correlation of the number of simulated feedback events to the composite measure. The data for all the possible pairs is shown in Figure 3c, with the correlation values of the two point and three-point algorithm outputs to the composite measure constituting the values along the X and Y axes respectively. Note that for almost all target pairs (49/55), the two-point method was better correlated to the composite measure than the three-point method, with 49 points falling below the identity line, and this difference was statistically significant (p<10^−4^, permutation test).

**Figure 3.**
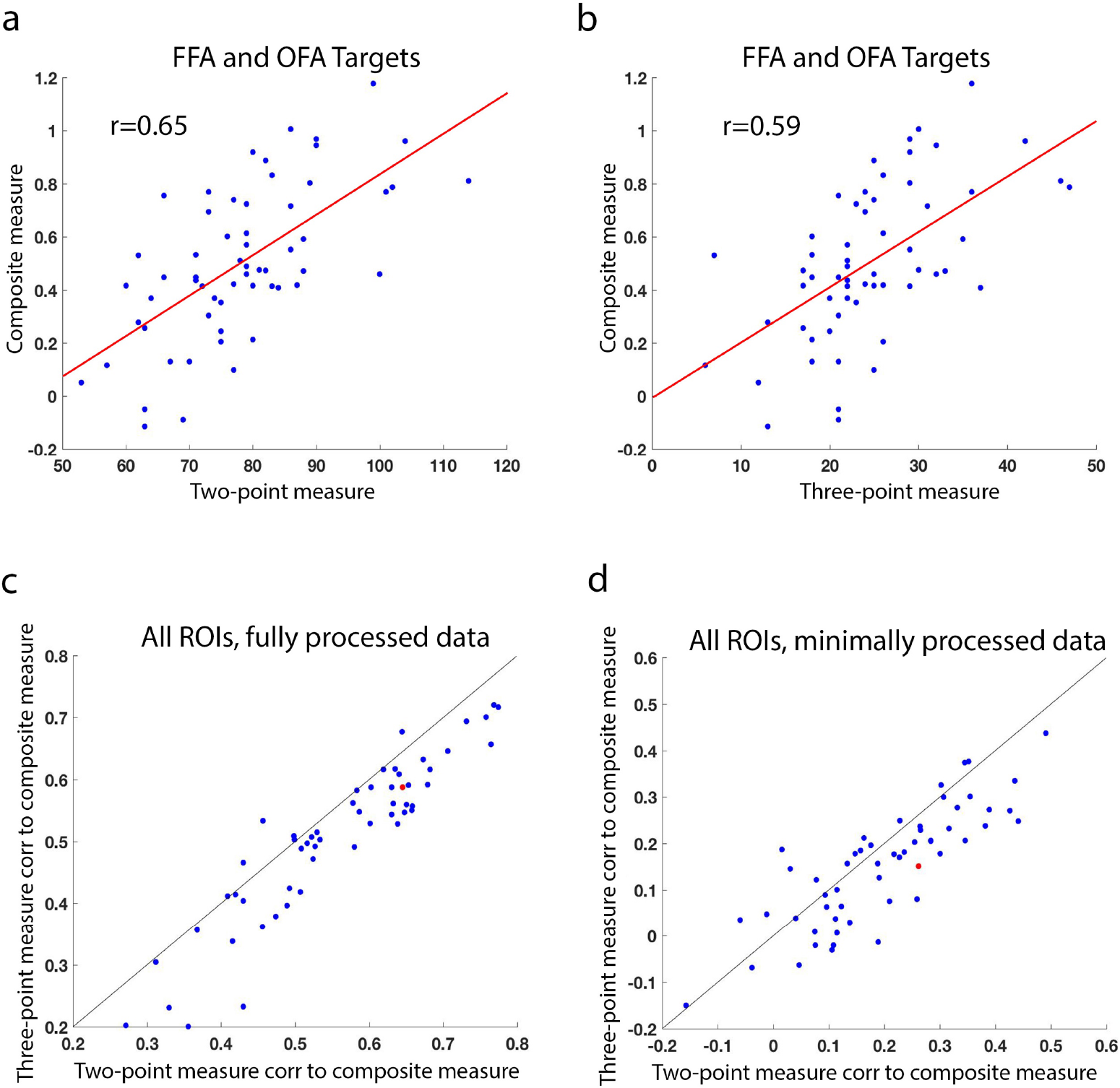
Number of time points. (a) The correlation between the output of the two-point algorithm on the fully processed data, with right FFA and right OFA as the targets on the x-axis (each dot represents the number of simulated feedback events for one participant, for one rest scan, 34 participants x 2 rest scans), and the composite measure (correlation of right FFA to right OFA, minus the average correlation of FFA to the control region and OFA to the control region) for each participant, for each rest scan. (b) Same as (a), but x-axis now shows the output of the algorithm using three time points instead of two (three-point algorithm). (c) The value for each of the 55 possible target pairs of the correlation between the two-point algorithm output and the composite measure (an example of which was shown in (a)), and the correlation between the three-point algorithm output and the composite measure (and example of which was shown in (b)). The right FFA/OFA pair shown in (a) and (b) is marked in red. Black line is the identity line, note that almost all dots (representing almost all target pairs) are to the right of the line, indicating the correlations are higher for the two-point algorithm. (d) Same as (c), but using the minimally processed data as input to the two-point and three-point algorithms right FFA/OFA target pair marked in red. Note that again correlation is higher for the two point method for most target pairs.

We then carried out the same analysis on the minimally pre-processed data, using only the preprocessing steps that would be available in our real-time scenario (3D motion correction, functional-anatomical co-registration). The noise levels in this analysis are higher because of the lack of sufficient pre-processing, so correlations are overall lower, but similarly to before, even with the increased noise, the two-point method still out-performed the three-point method for 73 percent of potential target pairs (40/55 pairs, Figure 3d). This difference was also assessed statistically through a permutation test, and was found significant (p<10^−4^).

### ROI size

Another important parameter to consider is ROI size. However, the optimal size for ROIs can be difficult to determine. Larger ROIs entail more averaging, which is beneficial for thermal noise reduction, but are also more likely to sample outside of the functionally relevant cortical region that was targeted. To determine the effects of ROI size, we first examined its influence on correlations between target regions, by measuring the full-series Pearson’s correlations between all possible 55 target pairs, in both the fully processed data (Figure 4a) and the minimally processed data (Figure 4b). In both cases, there was a linear increase in correlations between ROIs as ROI size increased, with differences between 4mm radius, 6mm radius, 8mm radius and 10mm radius ROIs all being significant (Figure 4c, p<10^−4^, permutation test). However, as our algorithm involves the interaction between the correlation of the targets with the correlation of the targets to the control (with reward given when the target regions are congruent but not congruent with the control region), an overall increase in correlation strength does not necessarily translate to better algorithm performance. We therefore next examined the effect of ROI size on the performance of the algorithm in the same manner as described above, by calculating the correlation of the simulated algorithm output for all 55 possible target pairs, using both the fully processed and the minimally processed data, to the full-series, fully processed composite measure (Figure 4d-f). For individual target pairs (Figure 4d-e), ROI size had quite a disparate effect, with larger ROIs increasing algorithm performance for some pairs, and decreasing it for others. When considering the average change across all possible target pairs, there was a significant increase in the correlation of algorithm output to the composite measure for the fully processed data between ROIs of size 10mm radius vs. all other ROI sizes (p<10^−4^, permutation test). The differences between all other ROI sizes were not statistically significant. For the minimally processed data, Only the 8mm radius ROIs were significantly better than others (p<10^−4^ for the difference between 8mm and the 4mm and 10mm ROIs, p=0.002 for the difference between 8mm and the 6mm ROIs, permutation test).

**Figure 4.**
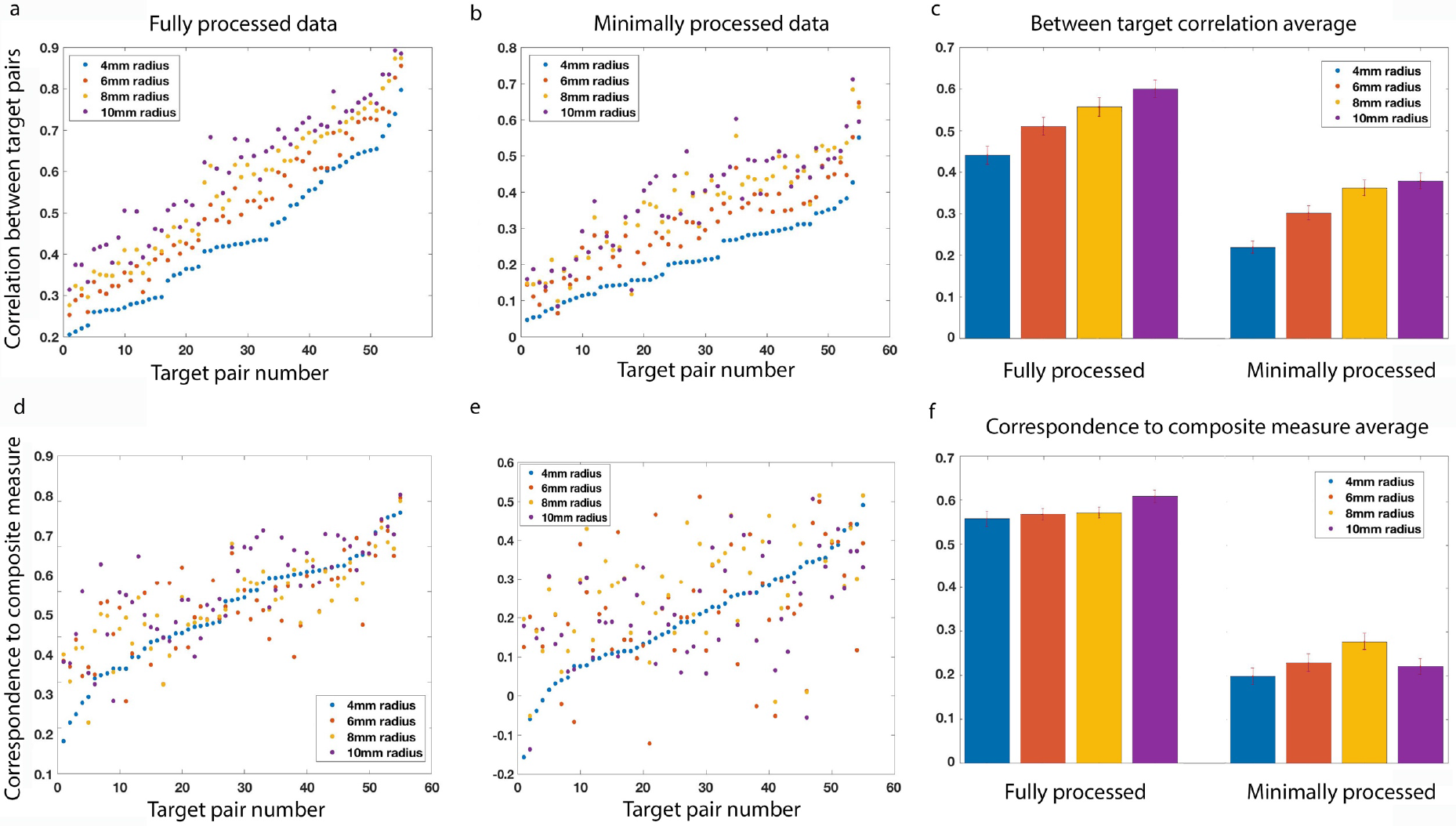
ROI size. Target-target full-series correlations for all 55 possible target pairs using 4mm radius ROIs (blue), 6mm radius ROIs (orange), 8mm radius ROIs (yellow) and 10mm radius ROIs (purple) on the fully processed data (a) and the minimally processed data (b). Target pairs sorted by target-target correlations of the 4mm ROIs. (c) Average target-target correlations for different ROI sizes for the fully processed data (left) and the minimally processed data (right). Error bar indicates standard error of the mean (SEM), colors same as (a) Correlations are significantly different between all the ROI sizes. (d) and (e) show the correspondence of the two-point algorithm to the composite measure for ROIs of size 4, 6, 8, and 10 mm radius for the fully processed and the minimally processed data, respectively. Target pairs sorted according to the 4mm ROI correlations. (f) Summary of (d) and (e) - average correlations of the algorithm to the composite measure for each of the ROI sizes, for the fully processed data (left) and the minimally processed data (right). Colors same as (a), error bar indicates SEM. Correlations are only significantly different between the 10mm ROI and the rest for the fully processed data, and between the 8mm ROI and the rest for the minimally processed data.

### Number of targets

We next tested whether adding additional targets to the algorithm would improve its correspondence to the composite measure. Two potential algorithm configurations were tested, using either three targets or four instead of two (see Materials and Methods). Again, we simulated the algorithm output for both the fully processed and the minimally processed data, and the results are shown in Figure 5. For each of the 55 possible target pairs, we plotted the correspondence to the composite measure for that target pair in red, with the boxplot showing the spread of correlations to the composite measure of all the possible combinations of 3 (top) and 4 (bottom) targets which include that pair. For the fully processed data, adding an additional target resulted in reduced correspondence to the composite measure for all possible target pairs / triads (Figure 5, top left). Adding two additional targets was also detrimental for all but 8 potential 4-target combinations (out of 3960 possible combinations) (Figure 5, bottom left). For the minimally processed data, adding targets had a more mixed effect, but additional targets only benefitted target pairs whose correspondence to the composite measure (without additional targets) was low to begin with (below r=0.23 for three targets, below r=0.3 for four targets). This happened in 13% of potential 3-target combinations, and 23% of potential 4-target combinations (Figure 5, right panels).

**Figure 5.**
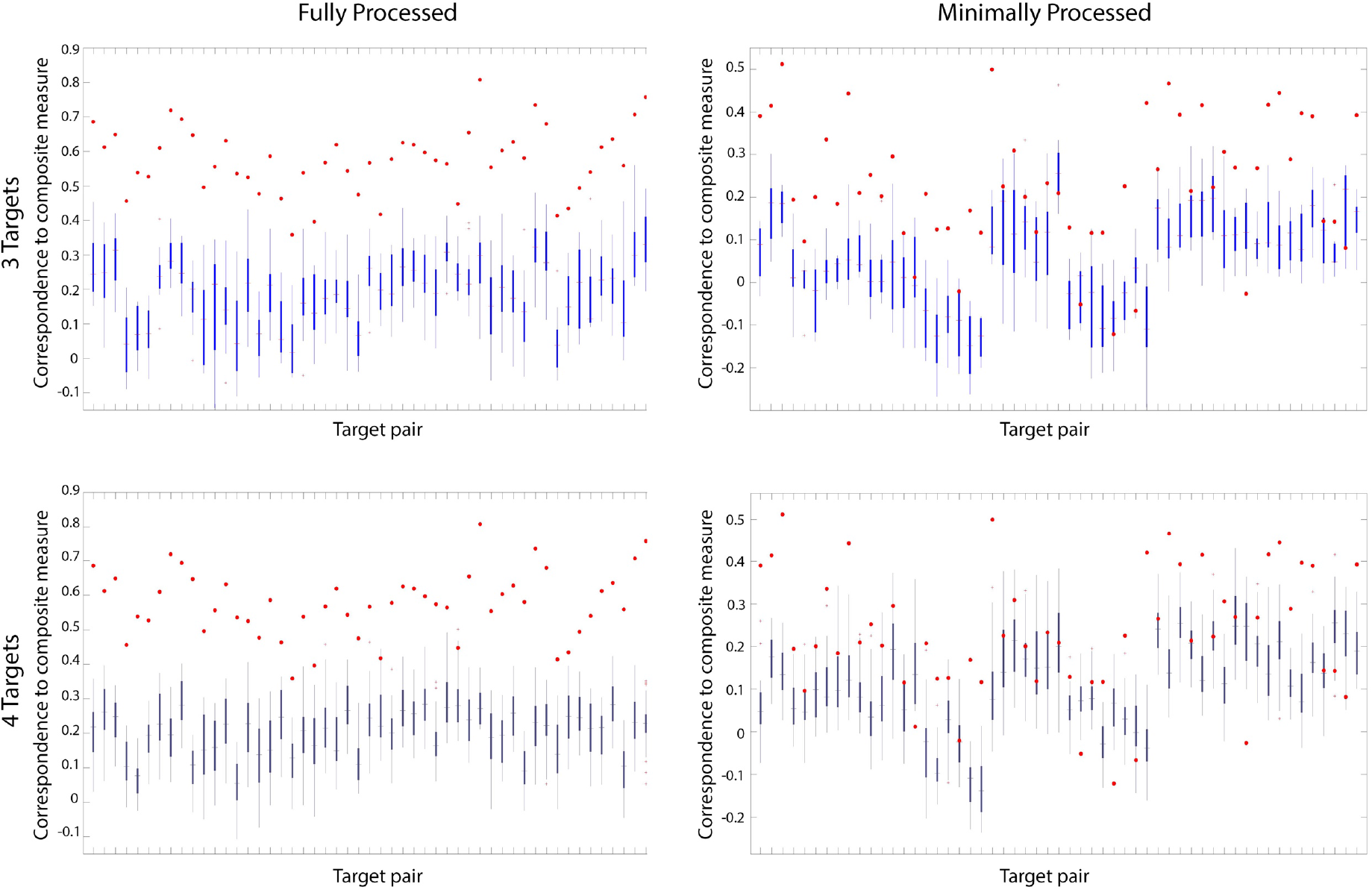
Number of targets. Top: for each target pair, comparing the correlation of the two-point algorithm to the composite measure using just these two targets (red dots) vs. all the possible correlations to the composite measure when adding a third target (box and whisker plots), for the fully processed data (left) and the minimally processed data (right). Bottom: same, but adding any other two targets to each possible target pair. Note that using just two targets gives better results, for all but the target pairs that perform very poorly to begin with.

### Optimally combined

Finally, we examined the effect of the optimal combination of the three echoes, a procedure which can be done online and thus a level of processing that could be available to the real-time algorithm.

As with the fully processed and the single-echo data, the OC data was sensitive to ROI size, with correlations between the 55 potential target pairs generally increasing with ROI size, but then declining when ROIs became too large (10mm, see Figure 6a-b). The differences between 4, 6, and 8mm ROIs were all significant (p<10^−4^, permutation test), whereas the 10mm ROIs had higher target-target correlations than the 4 and 6mm ROIs (p<, p=0.02, respectively), but lower correlations than the 8mm ROIs (p=0.046). However, for the real-time algorithm correspondence to the composite measure, the 4mm ROIs fared significantly better than all others (p<10^−4^, permutation test, Figure 6d-e). The OC target-target correlations for 4mm ROIs were slightly but significantly greater than the single echo target-target correlations (mean difference = 0.014, p<10^−4^, Figure 6c). The difference between OC and single echo in the correspondence of the algorithm to the composite measure however (again for the 4mm radius ROIs), was more pronounced (mean difference = 0.11, p<10^−4^, Figure 6f). For larger ROI sizes however, single-echo ROIs did better than the OC in terms of algorithm performance (p<0.01).

**Figure 6.**
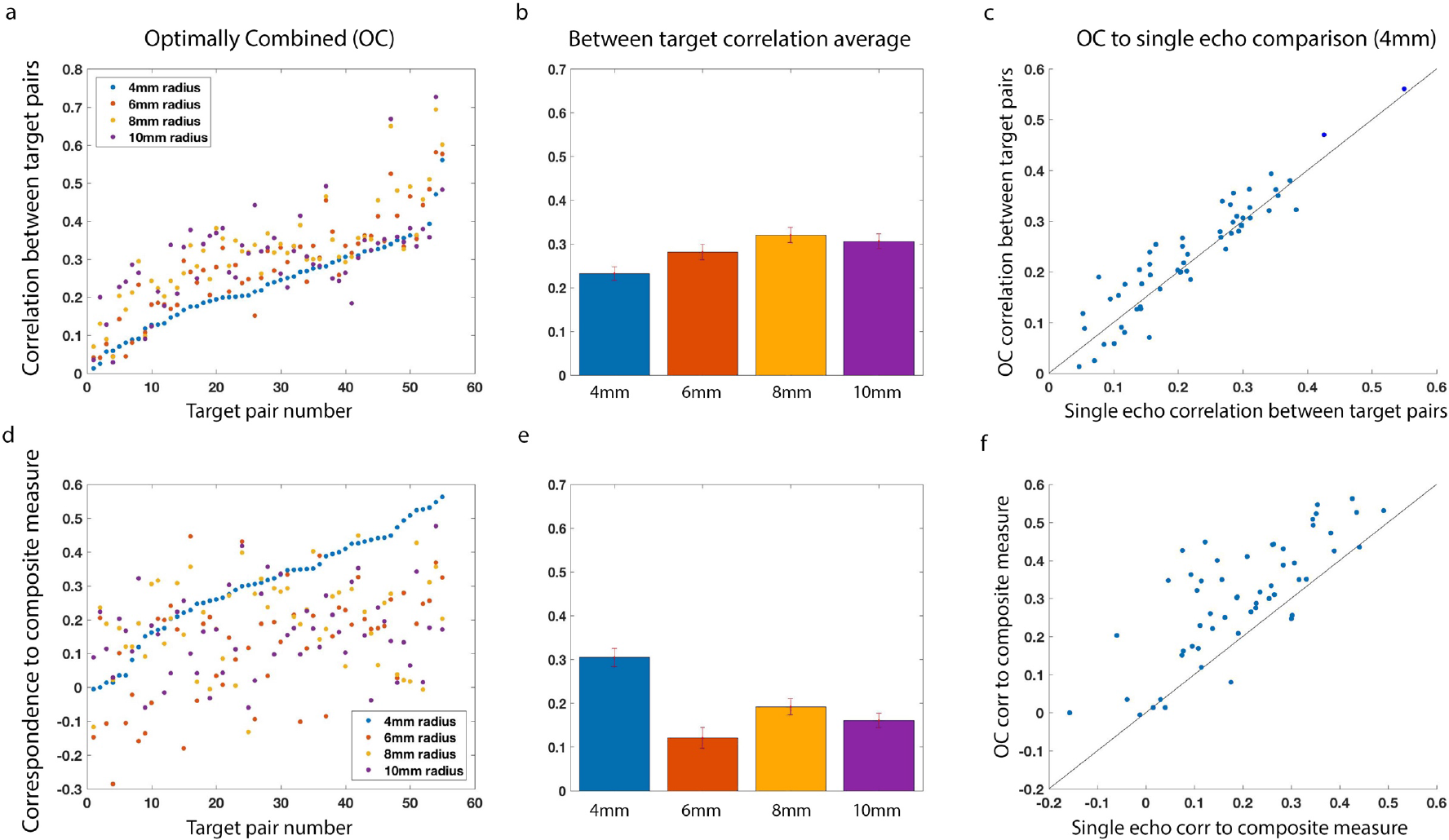
Effects of optimally combining echoes. (a) Target-target full-series correlations for all 55 possible target pairs using 4mm radius ROIs (blue), 6mm radius ROIs (orange), 8mm radius ROIs (yellow) and 10mm radius ROIs (purple) on the minimally processed data after optimal combination of the echoes (MP-OC). Target pairs sorted by target-target correlations of the 4mm ROIs. (b) Average target-target correlations for different ROI sizes for the MP-OC data. Error bar indicates standard error of the mean (SEM), colors same as (a). Correlations are significantly different between the 4mm, 6mm, and 8mm radius ROIs. 10mm radius ROI is significantly greater than the 4 and 6mm ROIs, but not the 8mm. (c) Comparing the target target correlations for the Minimally Processed Single Echo data (MP-SE) and the MP-OC data. There is a slight but significant difference in favor of the MP-OC data. (d) Correspondence of the two-point algorithm on the MP-OC data to the composite measure for ROIs of size 4, 6, 8, and 10 mm radius. Target pairs sorted according to the 4mm ROI correlations. (e) Summary of (d) - average correlations of the algorithm to the composite measure for each of the ROI sizes, for the MP-OC data. Colors same as (a), error bar indicates SEM. Correlations are only significantly greater for the 4mm ROI compared to the others. (f) Comparing the correspondence of the algorithm to the composite measure for the MP-SE data and the MP-OC data. There is a significant difference in favor of the MP-OC data.

### Target identity

Given the large degree of variance in algorithm performance for the different target pairs, we decided to test two possible hypotheses for the source of the at least a portion of that variance, the target-target correlations, and tSNR.

Target-target correlations were found to explain a large degree of the variance. This was especially true for the fully processed data, for which the correlation between the full-series, target-target Pearson’s correlation of each target pair, and the correlation of the algorithm output using that target pair to the composite measure was r=0.81 (measured across the 55 pairs, i. e. the correlation between the target-target correlations. For the minimally processed, single echo data this was lower, r=0.48, and for the minimally processed, optimally combined data, the correlation between these two measures was r=0.62. Correlation strength is influenced by the signal to noise ratio, as uncorrelated noise sitting on top of truly correlated signals can reduce the strength of correlation estimates. We therefore calculated the correlations between the target-target correlations for each target pair, and the average tSNR for all the voxels included in the two target ROIs for the fully processed and minimally processed data, with and without optimally combined echoes. The correlation of the tSNR to target-target correlations for the fully processed data was r=0.52, for the minimally processed data without optimally combined echoes r=0.66, and with optimally combined echoes r=0.61. This analysis suggested that although there is a significant correspondence between tSNR and target-target correlations at all levels of processing (p<10^−4^, permutation test), the cleaner the signal, the less the target-target correlations were affected by tSNR, therefore the variance in algorithm performance was potentially driven more by the target-target correlations irrespective of tSNR, than by tSNR itself. To test this, we calculated the partial correlations of the algorithm performance to target-target correlations, while co-varying out tSNR. For the fully processed data this correlation remained high (r=0.74), while for the single echo minimally processed data the correlation dropped to r=0.22, and for the OC minimally processed data it dropped to r=0.39, confirming the above hypothesis. This data is summarized in Figure 7.

**Figure 7.**
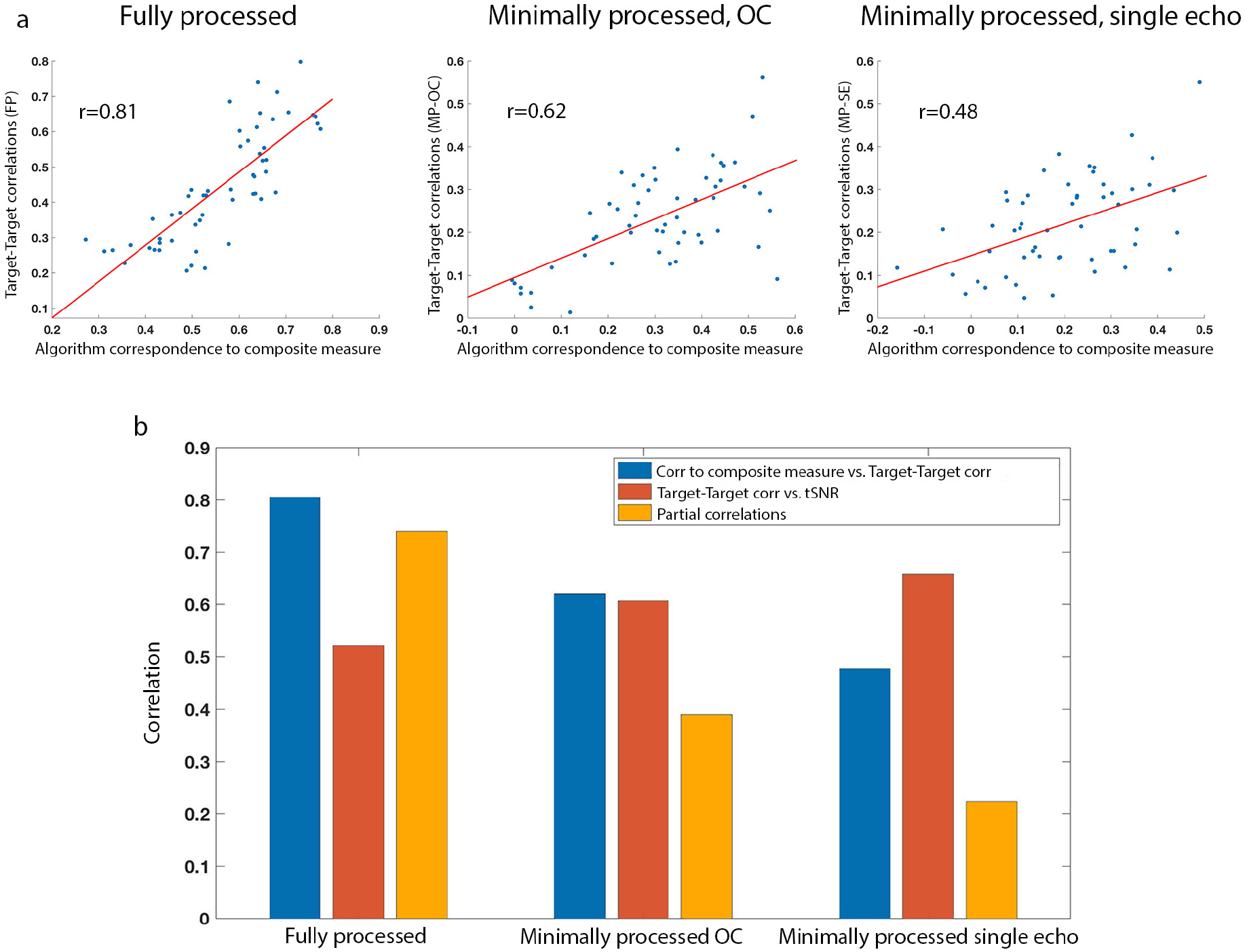
Explanation of variance. (a) Correlation of the correspondence of the two-point algorithm to the composite measure for each of the 55 target pairs (x-axis), to the target-target correlations (y-axis), for the fully processed (FP) data (left), the MP-OC data (center) and the MP-SE data (right). (b) Blue bars show the same correlation measure as (a), between the algorithm to the composite measure, and the target-target correlations, across all possible pairs, for FP (left), MP-OC, (center) and MP-SE (left). Orange bars show the correlations of target-target correlations to tSNR, and yellow bars show the partial correlations of the algorithm correspondence with the composite measure, to target-target correlations, after factoring out tSNR. Note that as data becomes better processed, the correspondence of the algorithm to the composite measure becomes more dependent on target-target correlations, but less dependent on tSNR.

### Comparing the algorithm to standard correlational approaches

Given the drop in performance for the algorithm using the minimally processed data, we wondered how this noisier data would affect calculation of Pearson’s correlations, which are more standardly used in connectivity-based feedback, for instance in sliding window approaches. We tested how these more traditional correlational methods would work for our training goal, to reinforce the correlations between the targets, and simultaneously decouple them from control. This would be most similar to calculation of the composite measure on the real-time data. For this we chose the least noisy data which could be made available to the feedback algorithm in an actual real-time environment, namely the OC minimally processed data. There are multiple different ways of calculating sliding window correlations, using different numbers of time points, and different averaging functions. Instead of calculating a version of the composite measure on all of these possible parameter combinations, we decided to simply calculate the composite measure on the full-series OC minimally processed data for each of the 55 possible target pairs, to get a more general estimate of how correlations differ between the fully processed and the noisier minimally processed data. Using the full time-series for this analysis would give us an upper bound on how well correlated sliding-window based algorithms can be to the fully processed composite measure. For each target pair, we calculated the composite measure for the OC minimally processed data and the fully processed data, for both rest scans, for each of the 34 participants, and then calculated the correlation between these two measures. An example of this for the FFA and OFA targets is shown in Figure 8a. As with the correspondence of the two-point algorithm output to the fully processed composite correlations, there was a large degree of variance in how well the composite measure on the OC minimally processed data corresponded to the composite measure on the fully processed data for different target pairs. To compare standard correlations to our two-point algorithm performance, we calculated the correspondence of the two-point algorithm (on the OC minimally processed data) to the fully processed composite measure for our 55 target pairs (Figure 8b, X-axis), and compared it to the correspondence of the composite measure on the OC minimally processed data to the composite measure on the fully processed data (Figure 8b, Y-axis). As can be seen from the figure, these were within the same range, though there were more target pairs for which the two-point algorithm was more highly correlated to the fully processed composite measure than the actual calculation of the composite measure on the OC minimally processed data. The difference between the two measures was significant (p=0.006, permutation test)

**Figure 8.**
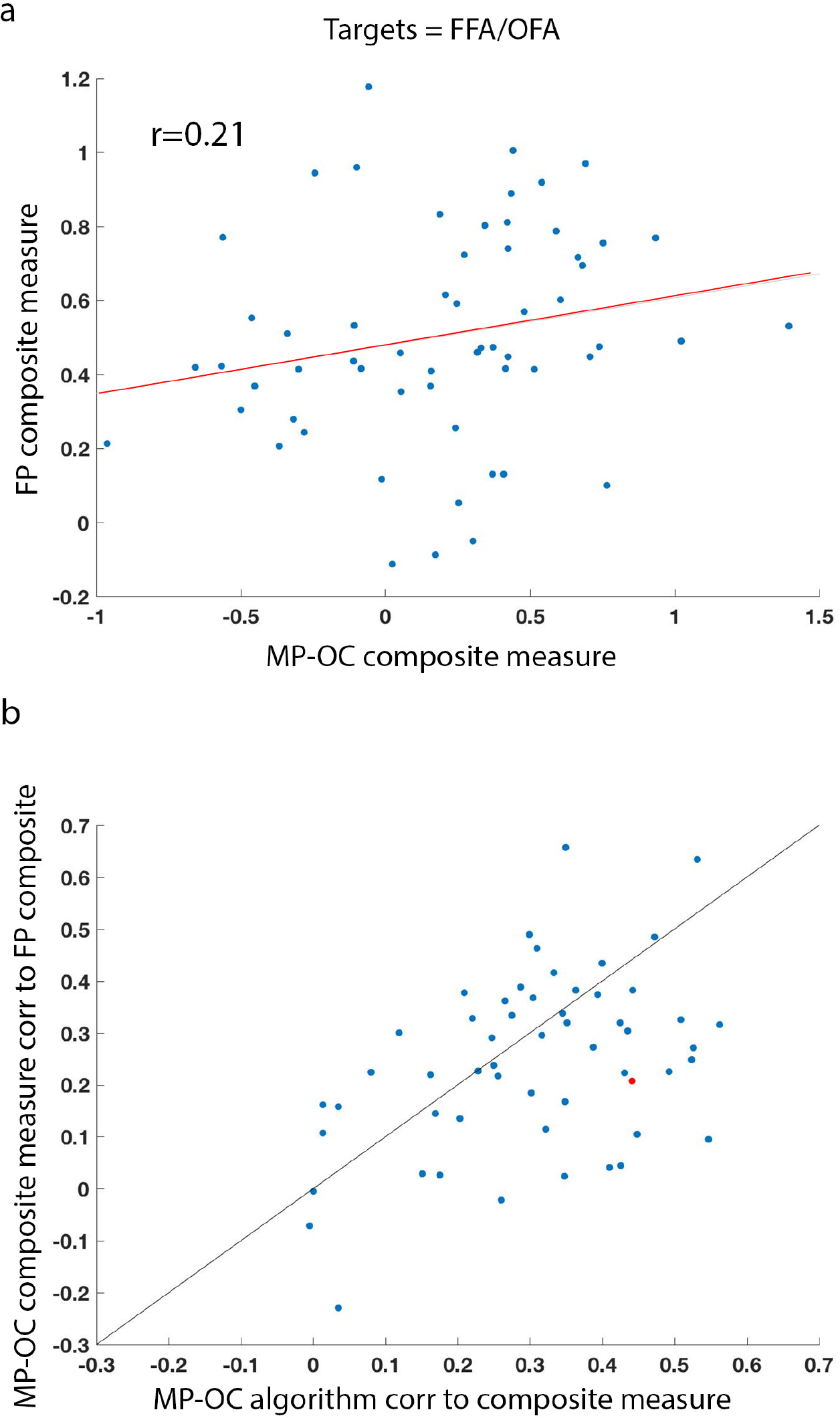
Comparing to more traditional correlation methods. (a) Comparing the composite measure calculated using right FFA and OFA as the targets, on the Minimally Processed Optimally Combined data (MP-OC, x-axis), and the fully processed data (FP, y-axis). Each dot represents the value of the composite measure for one rest scan, for one participant, on the MP-OC data (x) and the FP data (y). (b) Comparing the performance of the two point algorithm for the MP-OC data (correlation of the algorithm to the composite measure, x-axis), with the correlation of the actual calculation of the composite measure on the MP-OC data with the composite measure calculated on the FP data (y-axis), for all possible target pairs. Example from (a) of the FFA and OFA pair plotted in red. Note that correlations are generally within the same range, and for most possible target pairs there is an advantage for the two point algorithm over just calculating the composite measure on the MP-OC data using the full time series.

## Discussion

With the development of real-time fMRI neurofeedback, new and more complex algorithms are emerging, algorithms which go beyond simple measurement of amplitude and perform nontrivial calculations on the real-time signal. The framework suggested here aims to provide a method for more rigorously validating and optimizing these algorithms for best feedback results before putting them to use in an expensive real-time experiment, through offline testing using a previously collected dataset. This type of framework can benefit not only those seeking to do connectivity-based feedback, but will be useful for any feedback method in which there are parameters to be optimized, for instance MVPA classifiers or Dynamic Causal Modelling (comparing the results of the classifier on the fully processed vs. minimally processed data is an important sanity check, finding the optimal number of time points to feed into the classifier, optimal ROI size, etc.).

The simulations carried out here, using various possible parameters for the two-point method (target identity, number of time points, ROI size, number of targets), had two primary objectives. The first, to validate and optimize our correlation proxy using ideal, fully processed data. This is a first step towards creating such a proxy, as the fully processed data is our ground truth for the state of correlations in the brain. Whatever algorithm we propose to act as a proxy for correlations, must first be shown to be valid on the cleanest signals we have. For this reason, we included the Multi-Echo ICA cleaning step in the fully processed data, as this has been shown to increase statistical power and effect size estimations (35, 36).

The second step is validating and optimizing the algorithm on the minimally processed data, which is the data the algorithm will actually have access to in a real-time experiment. Though the algorithm performance (measured as the correlation of the algorithm output to the fully processed, full series composite measure) on the minimally processed data was predictably lower than on the fully processed data, reassuringly, the different parameters generally affected these correlations in much the same manner. Broadly speaking, it appeared that simpler was better. Fewer time points and fewer targets were better (Figures 3, 5), and larger ROIs increased algorithm performance only when ROIs were so large that they were unlikely to still be functionally congruent (10mm radius for the fully processed data, 8mm radius for the minimally processed data, Figure 4). For a discussion of functional ROI size within the face network see (37). In any case, it is important to find parameters which work well in both the fully processed and the minimally processed condition, so extremely large ROIs such as the 10mm radius ones which perform worse in the real-time conditions, are unlikely to be useful in this case. It may seem unintuitive that using more time points was detrimental to the algorithm’s performance, but it is important to note in this regard that this was true of the algorithm output on the fully processed data, as well as the minimally processed data, and is therefore unlikely to be purely due to the noise in the signal.

Acquiring multi-echo data has significant advantages in terms of post-processing, but to our knowledge, the advantages of the online optimal combination of the echoes to a real-time algorithm have not been tested. The results here suggest that at least for small ROIs (and probably also if using single voxels), the OC minimally processed data gives better results than the single echo data, though the advantage is lost for larger ROIs (Figure 6). This could be due to averaging in larger ROIs already reducing thermal noise to a similar extent to that achieved by OC, thus negating its advantage. Neither OC nor averaging over large ROIs address other sources of noise, such as motion artifacts or physiological noise, and these would still be present in the minimally processed data regardless of the chosen parameters. More advanced processing methods which can currently only be carried out offline on the full time-series, such as ME-ICA, or AFNI’s ANATICOR pipeline (38), would be needed to remove these.

Taken together, these results suggest that the best performance of this algorithm would be achieved by choosing only two targets (even though a wider network could well be functionally relevant), defining small (4mm radius) ROIs, using only two time points to determine feedback, and using online OC. We must however add a caveat, that all these analyses were carried out under specific acquisition parameters, and it is possible that with different acquisition parameters we would have found different results. For instance, a shorter TR might have resulted in more robust performance for the algorithm using three time points instead of two. Similarly, all the ROIs tested here were visual ROIs, and only one control region was tested. Different ROIs outside of visual cortex, or a different control ROI, might behave differently. Different neurofeedback objectives will impose different constraints regarding the location and size of target and control ROIs.

Throughout these analyses, what stood out perhaps even more than the effects of the various parameters we manipulated, was the great degree of variance among our 55 potential target pairs. This was apparent in all the parameters we compared, and was true for both the fully processed and the minimally processed data, with and without the optimal combination of echoes. In searching for the source of this variance, we discovered that much of it is explained by the target-target correlation within each target pair. This means that the identity of target ROIs is important for the success of the algorithm, and potential targets should be evaluated prior to beginning a new neurofeedback experiment, with baseline correlations between the targets being taken into account. Most connectivity-based training paradigms seek to increase correlations between regions with initially poor connectivity. The two-point algorithm might be less effective in training correlations between regions that are uncorrelated to begin with, though as long as some degree of initial correlation is present, training is possible (see Figure 7).

While tSNR does not appear to have a large effect on algorithm performance in the fully processed data (see partial correlations in Figure 7), it does greatly affect the minimally processed data, which is used for the actual real-time calculations. It would therefore seem prudent to steer away from signal-poor regions, such as ATL in our dataset, whenever possible. It should be noted that issues of noise affect not just this algorithm, but any calculation on real-time data. The two point algorithm actually performed overall better than a straightforward calculation of correlations on the OC minimally processed data, even using the full time series, and either way the two measures were within the same range (Figure 8). Note that this analysis only estimates the effects of noise on the correspondence of the online measures to our gold standard, offline calculation. It does not take into account other differences between the online algorithms, such as number of feedback events and reliance on history, which are likely to affect the induced training effects.

Beyond the technical points described here, the greatest challenge in crafting the best algorithm for providing the most accurate feedback, might be identifying the gold standard against which to compare it. In the demonstration of the two-point algorithm here, the goal was to train a small network of relative correlations between the target and control ROIs. Although our measure is instantaneous, theoretically allowing us to train dynamic connectivity, we are interested in training the static connectivity, which is the average connectivity throughout the entire scan. In our case, it is the static connectivity which was correlated to the behavior (26), and our gold standard was therefore the composite measure, comprised of the relative correlations between the targets and the control over the full time-series. When choosing the gold standard offline calculation for testing algorithm performance using this framework, it is imperative to consider the goal of the training. Had we been attempting to train dynamic connectivity, our gold standard would not have been the composite measure calculated over the full time-series, but something more relevant to such an alternative goal, such as the composite measure calculated with sliding windows.

Alternately, as the ultimate objective of the training is often to induce a change in behavior, and targets for training are generally chosen because of their correspondence to a behavioral measure, then an additional way to evaluate the validity and reliability of an algorithm would be through direct comparison to the relevant behavioral measure. The exact same general framework can be applied in this case as well, for instance comparing how correlated the algorithm output is to the behavioral measure using two time points vs. three time points. Comparing the algorithm output to the behavioral measure also provides an important sanity check. If there is no significant correlation between the two, then can the feedback induce the desired behavioral change? It is also possible for the two measures (one examining the network, such as correlations, and a second for behavior) to disagree on the optimal algorithm parameters.

ROI size is a likely example in this case. Although for the fully processed data, the correspondence of the algorithm to the composite measure increased with ROI size, it is doubtful that this would also be the case if comparing to a behavioral measure, given the extent of the functional averaging that must take place in an ROI of that size.

Overall, many of the optimal parameters for use with our two-point algorithm were unintuitive, illustrating the need for a framework such as this for testing out algorithms before applying them in an actual real-time experiment. Given the large degree of variance based on both scan parameters and target identity, it is recommended that an analysis using this framework be carried out in the design phase of any new experiment, using a dataset with similar parameters for testing.

## Acknowledgments

We are grateful to Dr. Alex Martin and Dr. Peter Bandettini for their continuous support and mentorship. We would like to thank Catherine Walsh, Kelsey Csumitta, and Jason Crutcher for help with data collection. This work was supported by the Intramural Research Program, National Institute of Mental Health (ZIAMH002920 and ZIAMH002783).

